# Tie2-Dependent Mechanisms Promote Leptomeningeal Collateral Remodeling and Reperfusion Following Stroke

**DOI:** 10.1101/2025.02.28.640890

**Authors:** Alexandra M. Kaloss, Caroline de Jager, Kennedie Lyles, Nathalie A. Groot, Jackie Zhu, Yu Lin, Hehuang Xie, John B. Matson, Michelle H. Theus

## Abstract

Leptomeningeal collaterals are distal pial arterial anastomotic vessels that provide an alternative route for redistributing cerebral blood flow following arterial obstruction, thereby limiting tissue damage. However, the regulatory mechanisms and strategies to enhance this adaptive response remain under investigation. This study explored the pharmacological effects of Tie2 receptor activation, using the peptide agonist Vasculotide, following permanent middle cerebral artery occlusion (pMCAO). Vasculotide improved collateral growth and remodeling, which correlated with reduced infarct volume, enhanced blood flow, and functional recovery within 24hrs post-pMCAO. In contrast, collateral growth was attenuated in Tie2 and EphA4/Tie2 double knockdown mice, while the loss of EphA4 increased Tie2 and Ang-1 expression and mimicked the positive effects of Vasculotide following stroke. Furthermore, bulk RNA sequencing of meningeal tissue identified key transcriptomic changes, including alterations in AJ-associated transcripts, such as *Krt5*, *Krt14*, and *Col17a1*, in the ipsilateral meninges of both endothelial cell-specific EphA4 knockout and Vasculotide-treated mice. Krt5 expression was found upregulated on meningeal arterial vascular network in injured KO mice, highlighting a potential new mediator of meningeal vascular remodeling. These findings illustrate that EphA4 and Tie2 play opposing roles in collateral remodeling, including the regulation of Krt5. Modulating their activity could potentially enhance the collateral response to stroke.

## Introduction

Ischemic stroke remains a significant global cause of mortality and morbidity. The loss of cerebral blood flow (CBF) following vascular obstruction leads to cell death and neurological impairments. It has been well documented that leptomeningeal anastomoses or pial collaterals can function to partially restore CBF to vulnerable neural tissue[1]. This retrograde perfusion is imperative for preserving the penumbral tissue and reducing tissue damage, making the extent of collateral vessels present a significant determinant of patient outcome following ischemic stroke[2]. Under healthy conditions, these specialized pre-existing arterioles are fed by two arteries, resulting in bidirectional blood flow. This, coupled with the high tortuosity and “non-physiological” angles of insertion, provides these vessels and their associated cells with a unique hemodynamic environment, as they are continuously exposed to disturbed flow [3]. Following an ischemic stroke, where one of the feeding arteries is blocked, these pial collateral vessels become exposed to unidirectional flow via the non-obstructed artery. This change in blood flow leads to an increase in fluid shear stress, the viscous drag force of the blood along the vessel wall, and triggers arteriogenesis, the remodeling process of these pial collaterals into conductance arteries[4].

The process of arteriogenesis requires endothelial and smooth muscle cell proliferation, immune cell recruitment, and degradation of the extracellular matrix to facilitate the outward expansion of the collateral vessel [5]. However, temporal cellular remodeling of the pial collaterals following ischemic stroke has not been fully established. Importantly, it remains unclear how early after occlusion these changes are evident and whether outward remodeling contributes to collateral size and redistribution of CBF. This knowledge gap in the acute remodeling capacity of pial collaterals could hinder the development of targeted therapeutics that enhance arteriogenesis or the vasodilatory capacity of the pial collateral vessels.

The receptor tyrosine kinase (RTK), Tie2, is essential in blood vessel formation and stabilization during development and in disease[6]. Its ligands, angiopoietin-1 (Ang-1) and angiopoietin-2 (Ang-2) bind with equal affinity but classically exert opposing effects, with Ang-1 acting as a Tie2 agonist and Ang-2 as a functional antagonist[7]. In a hindlimb ischemia model, transgenic mice overexpressing Ang-2 had smaller collateral artery size and reduced blood flow recovery[8]. Similarly, administration of recombinant Ang-2 reduced arteriole diameter and monocyte recruitment to the blood vessels[9]. However, the role of Ang-1 in pial collateral response remains less well-defined and represents an important knowledge gap. Moreover, prior studies have suggested that EphA4, another RTK, negatively regulates post-stroke collateral growth and may act as an upstream modulator of Tie2[10, 11].

This study demonstrates that a single dose of Vasculotide, an Ang-1 mimetic peptide, effectively enhances the pial collateral vessel response and promotes functional recovery. To further explore EphA4’s role as an upstream regulator of Tie2 signaling, we generated EC-specific EphA4 knockout (KO) mice. The KO mice exhibited increased Tie2 and Angiopoietin-1 (Ang-1) protein expression, which mirrored the enhanced collateral response observed in Vasculotide-treated mice. Uniquely we find Keratin 5 (Krt5), an epithelial structural protein, was commonly upregulated in Vasculotide-treated and KO-injured ipsilateral meningeal tissue that is expression on the arterial vasculature. These findings support the notion that the Tie2 axis is pivotal in dictating collateral response and can be a viable therapeutic target for augmenting collateral growth following an ischemic stroke.

## Results

### Tie2 agonist peptide, Vasculotide, protects against pMCAO and improves functional recovery

To interrogate the role of Tie2 in collateral response following pMCAO, Vasculotide, an angiopoietin-1 memetic peptide, was employed. Increased phosphorylation was seen in cultured endothelial cells (ECs) treated with Vasculotide compared to controls, demonstrating dose efficacy (Supplemental Figure 1). An intravenous bolus injection of Vasculotide was administered at 3μg/kg or 150μg/kg immediately following pMCAO. At both doses, Vasculotide decreased infarct volume compared to vehicle-treated mice (vehicle: 18.73 ± 1.58mm^3^; 3μg/kg: 9.09 ± 0.76 mm^3^; 150μg/kg: 11.42 ± 1.33 mm^3^) (Figure 1, A-B), which correlated with increased CBF recovery at 1-4 days post-stroke in 3μg/kg and at 1-2 days post-pMCAO in 150μg/kg (Figure 1C, 1D).

**Figure 1.**
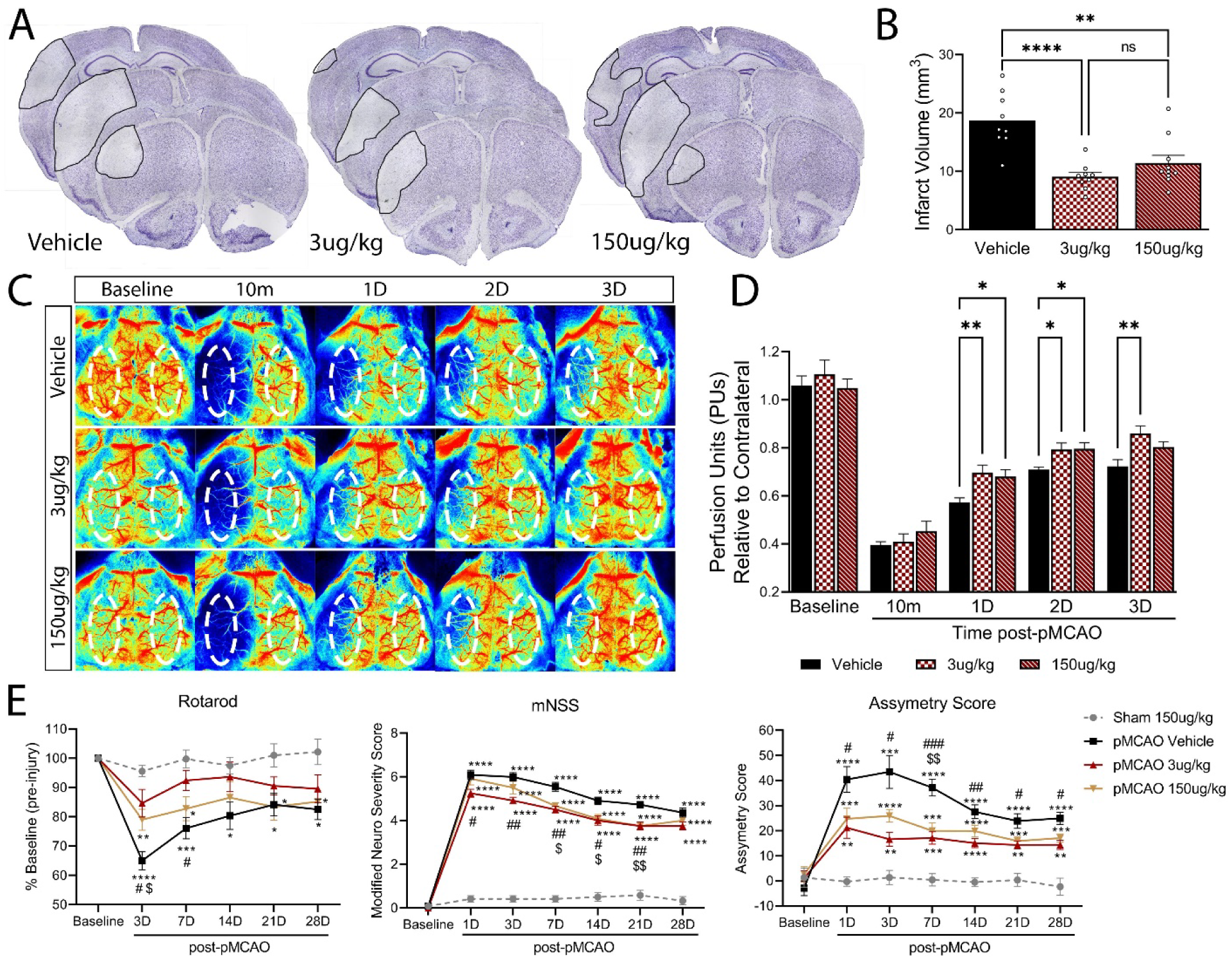
Angiopoetin-1 memetic peptide, Vasculotide (VT), confers neuroprotection and improves functional recovery in wildtype mice. (A) Representative images of Nissl-stained vessel painted sections 24hrs post-pMCAO from WT mice treated with saline (vehicle control), 3μg/kg-VT, or 150μg/kg-VT. (B) Quantified analysis of infarct volume shows Vasculotide confers significant neuroprotection. n=9-10. (C) Representative laser speckle contrast images of CBF in vehicle- and Vasculotide-treated mice. (D) Quantified CBF analysis pre- and post-pMCAO. n=12 (E) Behavioral analysis using Rotarod (left), modified neurological severity score (center), and adhesive tape test (right) showed improved performance in Vasculotide treated mice, notably the 3μg/kg-VT group, compared to vehicle controls. n=11-12. Analysis was completed using ordinary one-way ANOVA (B) or two-way ANOVA (D-E) with Tukey’s post hoc analysis. Dotted oval in E = standardized ROI used to measure CBF. * = compared to Sham 150μg/kg-VT mice. # = pMCAO 3μg/kg-VT vs pMCAO Vehicle mice. $ = pMCAO 150μg/kg-VT vs pMCAO Vehicle mice. ^*,#,$^p<0.05, ^**,##,$$^p<0.01, ^***,###^p<0.001, ****p<0.0001.

Moreover, mice receiving 3μg/kg Vasculotide showed reduced motor deficits on the rotarod at 3- (vehicle: 65.00 ± 2.94% vs. 3g/kg: 84.67 ± 4.72%) and 7-days post-pMCAO (vehicle: 76.09 ± 3.48% vs. 3μg/kg: 92.42 ± 3.46%). In comparison, 150μg/kg Vasculotide showed reduced deficits only 3 days post-stroke (79.00 ± 60%). Likewise, treatment with Vasculotide reduced sensorimotor deficits assessed by Modified Neurological Severity Scoring (mNSS) and Asymmetry Score, most notably in the 3μg/kg Vasculotide group (Figure 1E). These findings suggest Tie2 activation is neuroprotective, and a single dose of Vasculotide can provide long-term functional improvements.

### Vasculotide enhances pial collateral response following stroke

Pial collateral vessel growth and remodeling support restoration of CBF and functional recovery after ischemic stroke. To determine whether Vasculotide-treated mice display reduced deficits through alterations in the acute collateral response, we performed vessel painting at 1-, 4-, and 28 days post-pMCAO. Mice receiving Vasculotide (3μg/kg or 150μg/kg) exhibited significantly larger ipsilateral MCA-ACA collaterals 1-day post-pMCAO compared to vehicle controls (Vehicle: 29.60 ± 0.80 μm; 3μg/kg: 37.16 ± 0.64 μm; 150μg/kg: 34.94 ± 0.98 μm). At 4-days post-pMCAO, the mice receiving 3μg/kg VT retained enlarged collateral sizes compared to both vehicle controls and the 150μg/kg-VT group (Vehicle: 44.41 ± 1.31 μm; 3μg/kg: 52.93 ± 2.73 μm; 150μg/kg: 44.70 ± 2.45 μm). By 28 days post-stroke, no significant difference in ipsilateral collateral size was appreciable (Vehicle: 40.91 ± 1.57 μm; 3μg/kg: 42.57 ± 2.22 μm; 150μg/kg: 39.91 ± 1.80 μm) (Figure 2A, 2C). Additionally, no changes were observed in the MCA-ACA collateral vessel of the contralateral hemisphere across treatment groups or time points (Supplemental Figure 2A). Collateral enhancement by Vasculotide was also reflected in the distribution of collaterals in the ipsilateral hemisphere, with the 3μg/kg group having significantly more collaterals over 40μm at 1- and 4 days compared to vehicle controls (Supplemental Figure 2B-2D). Across all treatment groups and time points, the ipsilateral MCA-PCA collateral size markedly increased compared to the contralateral hemisphere. However, no differences were noted between the ipsilateral size of the treatment groups (Supplemental Figure 2E).

**Figure 2.**
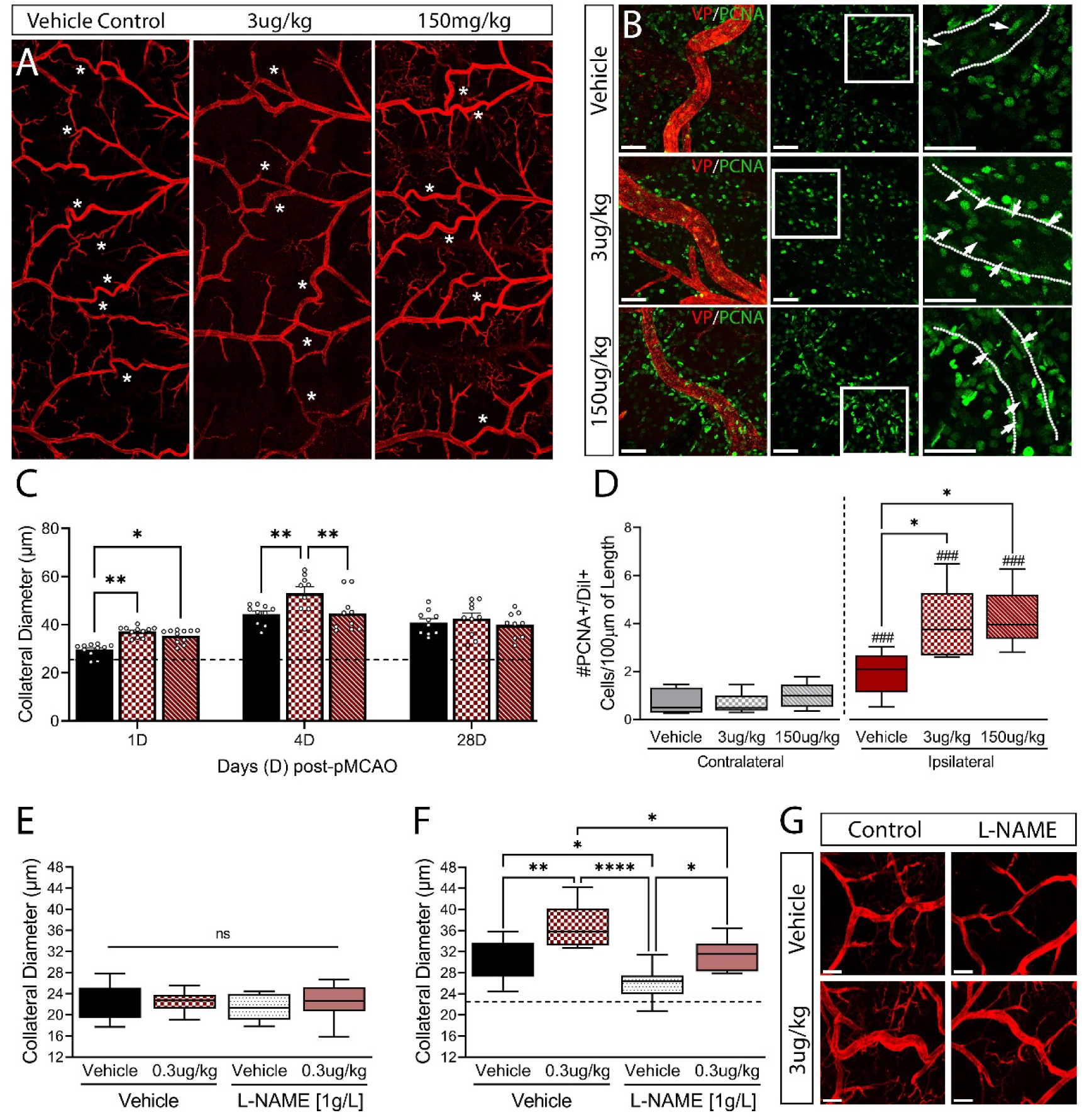
Angiopoetin-1 memetic peptide, Vasculotide (VT), leads to increased collateral growth and endothelial cell proliferation. (A) Representative tiled 4X confocal images of vessel painted brains 1 day post-pMCAO in mice receiving vehicle, 3μg/kg-VT, or 150μg/kg-VT. (B) Representative 40X images of PCNA stained MCA-ACA pial collateral vessels at 1-day post-pMCAO from mice receiving vehicle, 3μg/kg-VT, or 150μg/kg-VT. (C) Analysis of MCA-ACA pial collaterals shows a significant increase in collateral diameter 1- and 4-days post-injury following 3μg/kg VT treatment. n=9-11. (D) Quantification of PCNA-positive ECs reveals Vasculotide treatment at either dose increases proliferation of ECs at 1-day post-injury. n=5. (E) L-NAME, a nitric oxide inhibitor, was administered in drinking water for 24 hours post-pMCAO. No significant changes were observed in contralateral MCA-ACA collaterals in response to either drinking water or 3μg/kg-VT treatment at 1-day post-pMCAO. (F) In the ipsilateral hemisphere, L-NAME significantly decreased MCA-ACA collateral size in both vehicle- and VT-treated mice compared to control drinking water; however, collaterals in 3μg/kg VT-treated mice remained larger than in vehicle-treated mice. Analysis was completed using ordinary one-way ANOVA (D) or two way ANOVA (A, E-F) with Tukey’s post hoc analysis. Scale Bar = 1mm (A-C), 50um (B), or 100um (G). Dashed line in panels C and F = average contralateral collateral diameter. #=compared to respective contralateral hemisphere. ^*,#^p<0.05, ^**,##^p<0.01, ^***,###^p<0.001, ****p<0.0001.

Enlargement of the pial collateral vessels into conductance arteries occurs due to the expansion of the endothelium through cellular proliferation and immune cell recruitment. Confocal imaging and quantification of proliferating cell nuclear antigen (PNCA) labeled, Dil-positive cells in the collateral wall showed a significant increase at 1 day in Vasculotide-treated mice, indicating increased numbers of proliferating ECs in MCA-ACA collaterals compared to vehicle-treated mice (vehicle: 1.94 ± 0.41; 3μg/kg: 3.92 ± 0.70; 150μg/kg: 4.22 ± 0.56) (Figure 2B, 2D).

Next, we investigated whether nitric oxide-induced vasodilation mediates the collateral effects in Vasculotide-treated mice. Nitric oxide (NO) is a free radical that is a potent vasodilator released after ischemic stroke that may impact collateral circulation [12]. To test this, L-NAME – a NO synthase inhibitor – was given to Vasculotide and vehicle-treated mice via drinking water at 1g/L for 1 day post-pMCAO. L-NAME treatment did not alter the collateral size in the contralateral hemisphere, indicating minimal NO-induced vasodilation on the uninjured hemisphere (Control water: 22.18 ± 1.038 m vs. 22.46 ± 0.63 m; L-NAME: 21.26 ± 0.75 m vs. 22.53 ± 1.09 μm) (Figure 2E). Compared to control water, L-NAME drinking water significantly decreased ipsilateral collateral size in the vehicle and 3g/kg-VT treated mice on the ipsilateral hemisphere. Interestingly, the 3μg/kg-VT mice on L-NAME drinking water retained significantly larger MCA-ACA connecting collaterals compared to vehicle mice on L-NAME water, indicating the enhanced collateral size seen with Vasculotide treatment is not solely due to NO-driven vasodilation (Control water: 30.61 ± 1.17 μm vs 36.50 ± 1.40 μm; L-NAME: 25.93 ± 0.90 μm vs 31.46 ± 0.98 μm) (Figure 2F-2G). Taken together, these findings demonstrate that Tie2 stimulation may be a crucial target for therapeutic intervention following ischemic stroke and that Tie2 enhancement of pial collaterals triggers arteriogenesis and vasodilation.

### RNA transcriptomic profile of the meninges reveals common alterations following Vasculotide treatment after pMCAO

To improve our understanding of the underlying changes in the meningeal environment that may support collateral growth, the pial surface was isolated at 1-day post-pMCAO (Figure 3A), and bulk RNA sequencing was performed in Vehicle or 3μg/kg Vasculotide treated mice. Mice treated with Vasculotide had the highest number of unique, differentially expressed genes, with 466 upregulated and 714 downregulated, compared to vehicle-treated mice (Figure 3A). Top ten genes between ipsilateral and contralateral meninges in Vehicle or Vasculotide conditions (Figure 3B), as well as between Vasculotide and Vehicle ipsilateral or contralateral tissue (Figure 3C) further highlights Krt5 as highly expressed following Tie2 stimulation. GO terms from genes induced in the ipsilateral meninges following Vasculotide treatment included meiotic cell cycle, mitotic and nuclear division, organic anion transport, cell chemotaxis, leukocyte migration, and cell adhesion (Figure 3D). Ipsilateral vs Contralateral, Vehicle treated GO terms included regulation of membrane potential, neurotransmitter transport, exocytosis, leukocyte migration, regulation of inflammatory response, and cell junction assembly (Figure 3E) compared to Vasculotide GO terms which included cognition, ribonucleoprotein complex biogenesis, regulation of developmental growth and epithelial proliferation (Figure 3F).

**Figure 3.**
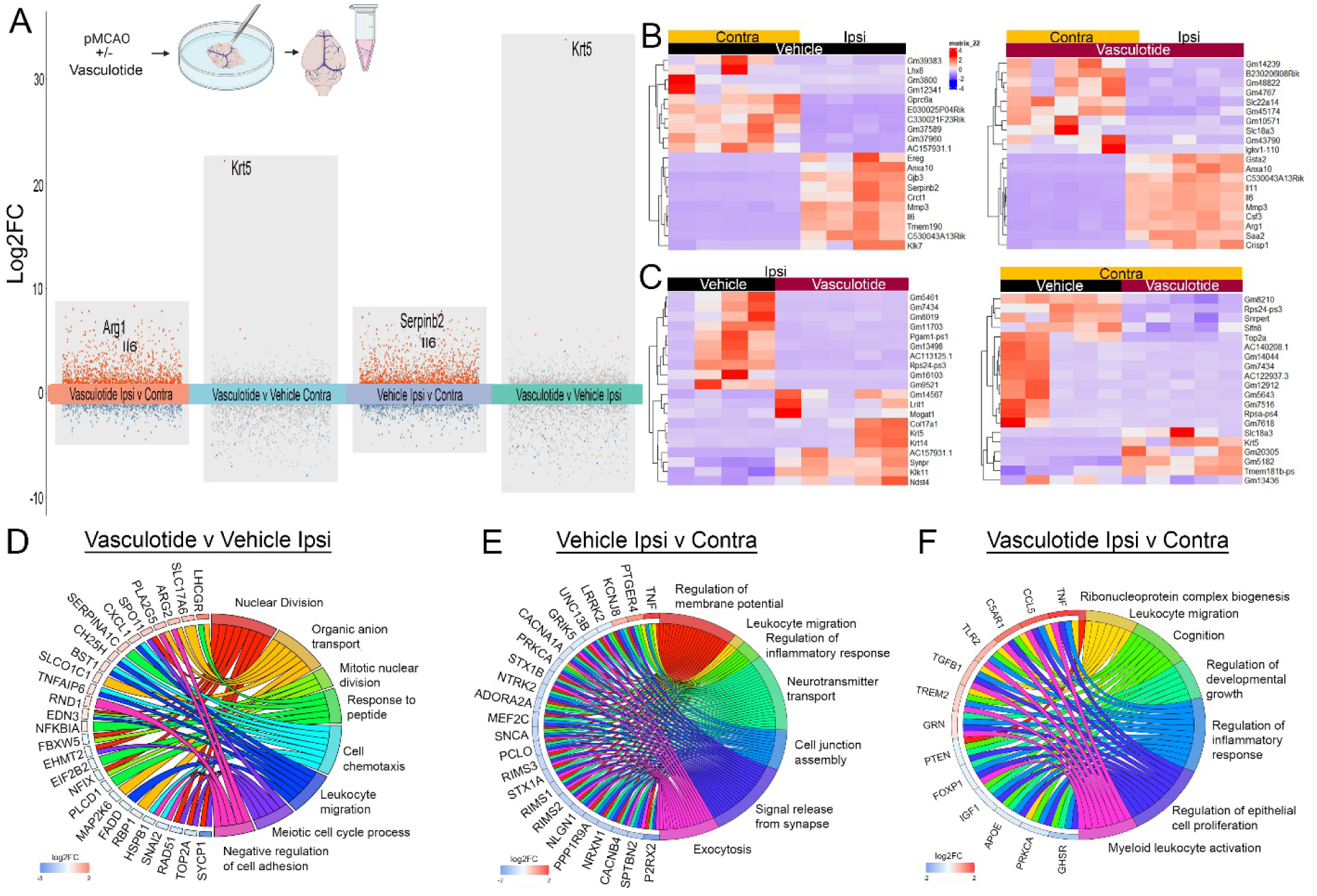
RNA sequencing of meningeal isolates displays altered transcriptomics in Vasculotide-treated mice. (A) Schematic of meningeal removal and RNA isolation. Transformed plot of bulk RNA sequencing from Vehicle and Vasculotide-treated meninges at 24hrs post-pMCAO. (B-C) Heatmap of top 10 upregulated and downregulated genes across conditions. (D-F) Chord plot of genes and GO terms between Vehicle and Vasculotide-treated ipsilateral and contralateral meninges.

### Endothelial EphA4 restricts Tie2 signaling to limit collateral growth and remodeling

Evidence from previous studies has implicated EphA4 as an upstream regulator of Tie2 activation post-pMCAO. To further elucidate the interplay between EphA4 and Tie2, we employed EC-specific EphA4 knockout mice (*EphA4^fl/fl^/VECadherin-CreERT2*; EC-specific KO) and wildtype controls (*EphA4^fl/fl^*; WT). Analysis of protein expression at 1-day post-sham and pMCAO revealed a significant increase in Tie2 (Sham: 0.39 ± 0.03 vs. 0.74 ± 0.06; pMCAO: 0.36 ± 0.05 vs. 0.57 ± 0.05, relative to b-actin) and angiopoietin-1 expression in the cortex of EC-specific EphA4 KO mice compared to WT controls (Sham: 0.13 ± 0.01 vs 0.33 ± 0.06; pMCAO: 0.13 ± 0.02 vs 0.38 ± 0.07, relative to b-actin). No differences were noted in Ang2 expression. These results indicate that EphA4 may suppress Tie2 expression (Figure 4A, Supplemental Figure 4A-4C).

**Figure 4.**
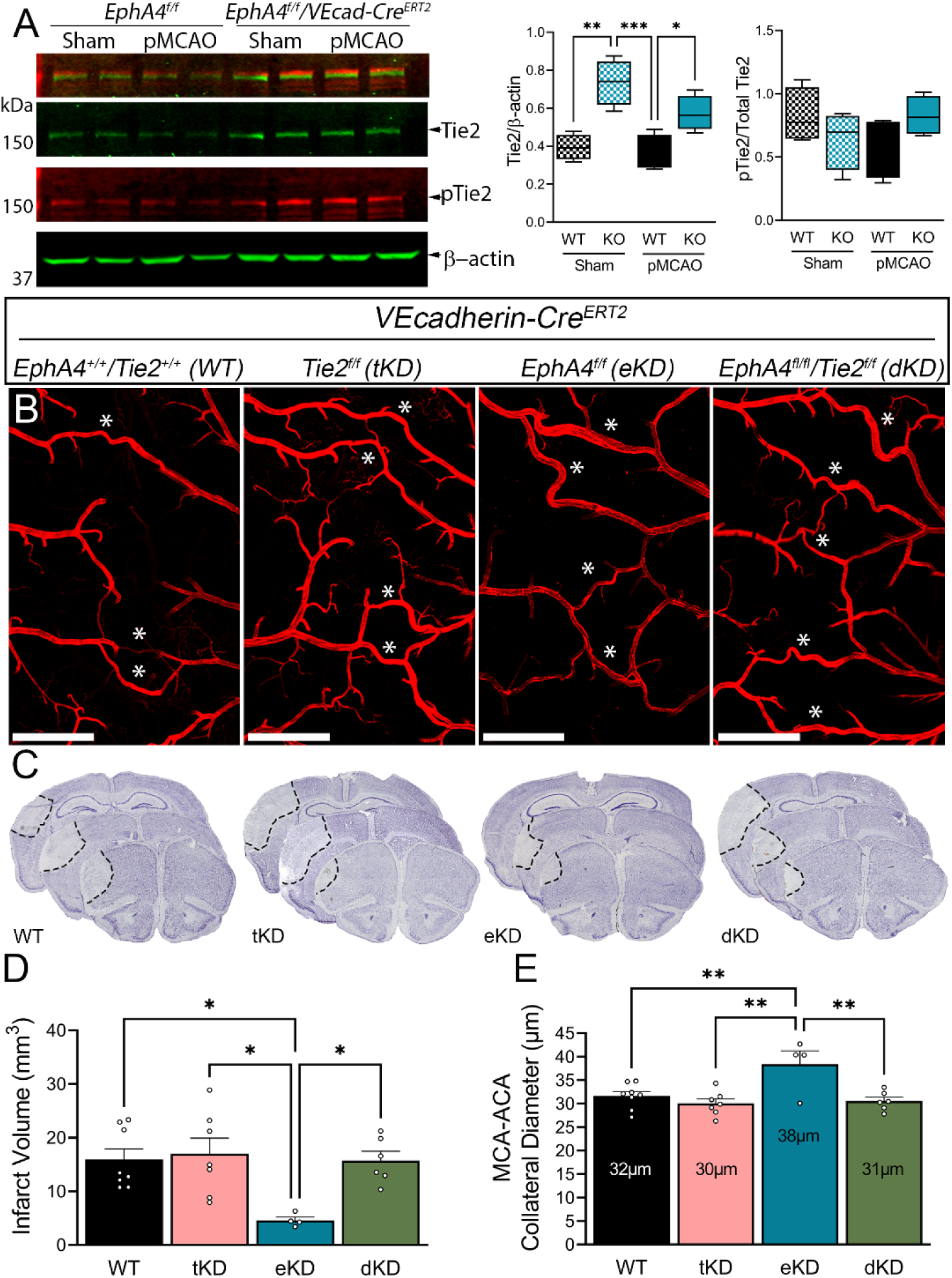
EphA4 serves as an upstream regulator of Tie2 and its loss increases Tie2 expression. (A) Western blot analysis of cortex samples 24hrs post-pMCAO from WT and EphA4 KO mice. Densitometric analysis shows increased Tie2 levels but no change in the proportion of pTie2 to total Tie2. n=4. (B) Representative 4X tiled confocal images of the ipsilateral MCA-ACA pial collaterals in WT, tKD, eKD, and dKD vessel painted brains. (C) Representative images of infarct volume from cryo-sectioned vessel painted brains. (D) Assessment of infarct volume shows a decrease in tissue damage only in the eKD mouse line, no changes were noted in the tKD or dKD mice compared to WT controls. (E) Similarly, compared to WT controls, only the eKD mice exhibited increased MCA-ACA collateral diameters. n=4-7. Analysis was done using a one-way ANOVA with Tukey’s post-hoc analysis. Scale bars = 1mm. *p<0.05, **p<0.01, ***p<0.001.

To examine whether EphA4 negatively regulates pial collateral remodeling by suppressing Tie2 signaling, we performed pMCAO on *Tie2^f/f^/VEcad-CreERT2 (tKD), EphA4^f/f^/VEcad-CreERT2 (eKD),* and *EphA4^f/f^/Tie2^f/f^/VEcad-CreERT2 (dKD)* mice 2-weeks post-tamoxifen injections. Tie2 deletion was confirmed by immunostaining (Supplementary Figure 3, D-F). While the loss of Tie2 signaling on ECs did not influence infarct volume or collateral diameter compared to WT, double EC knockdown of EphA4 and Tie2 attenuated the positive effects observed in single EphA4 knockout mice (Figure 4B-4E). These findings confirm that EphA4 primarily affects collateral growth by restricting Tie2 signaling.

Lastly, due to Eph/ephrin bi-directional signaling [13], we investigated the directional effects of EC-specific loss of EphA4 to further understand its control over Tie2. Clustered (cl) EphA4-Fc ectodomain was intravenously injected following pMCAO to restore reverse signaling while maintaining forward endothelial deletion. Compared to Fc-control, no difference in collateral diameter or number was observed at 1 day in clEphA4-Fc-treated KO mice (Supplemental Figure 4). This supports the role of EphA4 forward signaling in restricting collateral growth.

### Loss of EC-specific EphA4 confers neuroprotection and improves functional recovery following pMCAO

To assess whether EC-specific EphA4 regulates ischemic stroke outcome, we employed EC-specific knockout mice (*EphA4^fl/fl^/VECadherin-CreERT2*; EC-specific KO) and wildtype controls (*EphA4^fl/fl^*; WT). Findings demonstrate a reduction in infarct volume in KO (14.31 ± 2.52 mm^3^) compared to WT mice (24.04 ± 1.69 mm^3^) at 1-day post-pMCAO injury (Figure 4A). This correlated with improved CBF perfusion in KO compared to WT control mice at 1-4 days post-injury (Figure 5B). KO mice also displayed marked improvements in behavioral recovery as demonstrated by Rotarod assessment of motor function at 3 (67.31 ± 3.45% vs. 78.77 ± 2.12%) and 7 days post-injury (76.84 ± 2.76% vs. 88.77 ± 2.89%), adhesive tape removal using asymmetry score of sensorimotor function, and modified neurological severity scores at 3 (5.69 ± 0.29 vs. 4.28 ± 0.21) and 7 days post-injury (5.08 ± 0.18 vs. 4.31 ± 0.17) (Supplementary Figure 4). These findings suggest that EC-specific EphA4 exacerbates tissue loss, impedes CBF recovery, and impairs functional recovery after injury.

**Figure 5.**
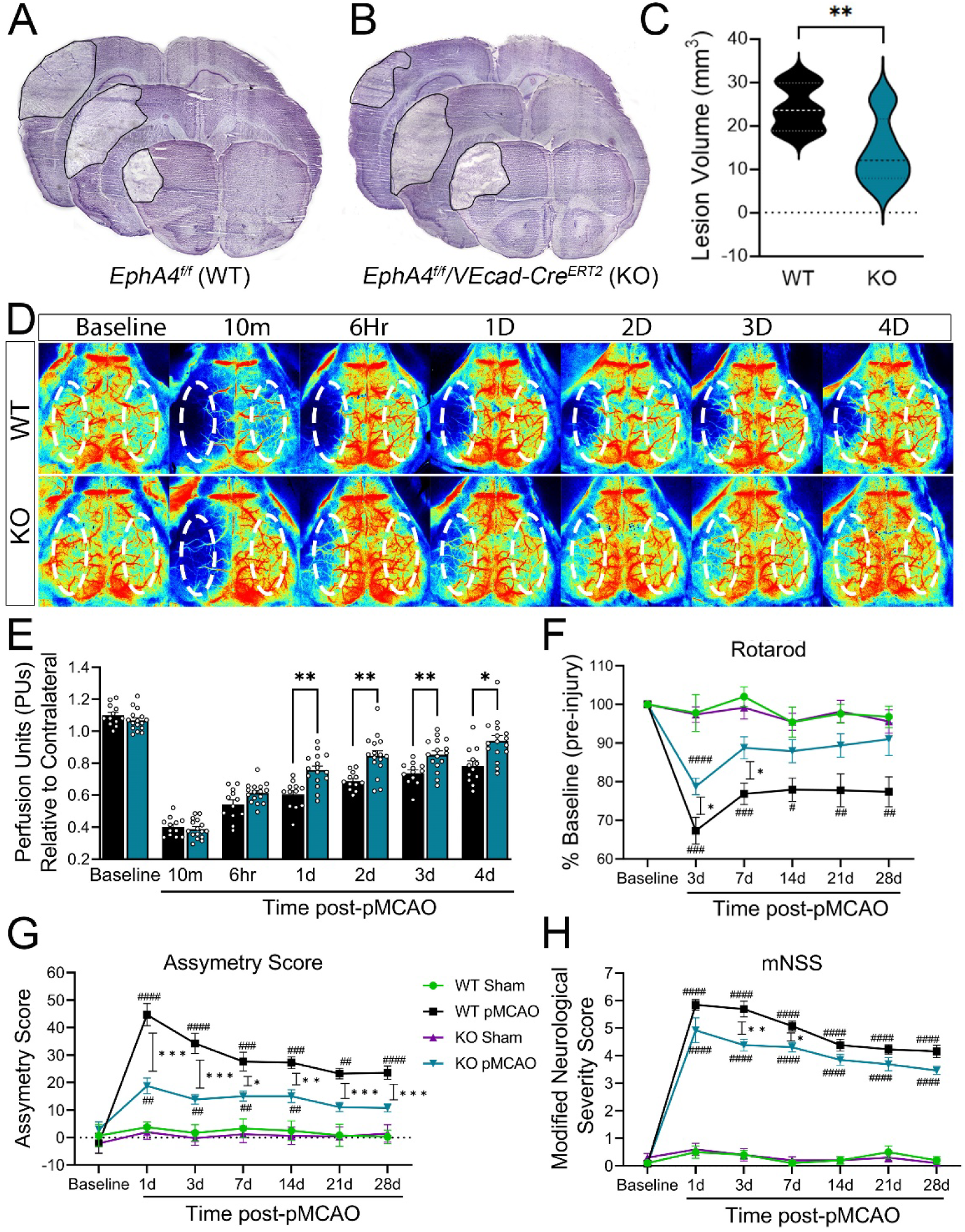
EC-specific EphA4 KO improves collateral growth and is partially attenuated with L-NAME treatment after pMCAO. (A-B) Representative fresh frozen Nissl-stained images of WT and EphA4 KO brains. (C) Quantified t-test analysis of infarct volume showed neuroprotection in the KO mice at 24hrs post-pMCAO. n=9 (D) Representative laser speckle contrast images pre-and post-pMCAO of WT and KO mice. (E) Quantification of CBF shows increased perfusion in KO mice at 1-4 days post-pMCAO. n=12-16. (F) Assessment of functional recovery using Rotarod, (G) adhesive tape test, and (H) modified neurological severity score reveal improved performance by KO mice compared to WT animals post-pMCAO. N=10-13. Analysis of E-H was done using a two-way ANOVA with Sidak’s (E) or Tukey’s (F-H) post-hoc analysis. Dotted oval in panel B=standardized ROI used to measure CBF. *P<0.05; **P<0.01; ***P<0.001 compared to WT mice. #P<0.05; ##P<0.01; ###P<0.001; ####P<0.0001 compared to respective sham groups.

### Loss of EC-specific EphA4 augments acute pial collateral growth after pMCAO

To determine if the increased Ang1 and Tie2 seen in EphA4 KO mice would recapitulate the findings of Vasculotide treatment, vessel painting and collateral analysis were performed. While no difference was observed between un-injured WT and KO collateral number or diameter (Supplementary Figure 5A-5B), KO mice showed a significant increase in the MCA-ACA ipsilateral collateral diameter (32.41 ± 0.77 μm) compared to WT (27.31 ± 0.58 μm) as early at 4.5hrs post-pMCAO, which was continued through 6hrs (WT: 29.41 ± 0.60 μm; KO: 33.70 ± 0.61 μm), 1-day (WT: 31.14 ± 0.84 μm; KO: 36.73 ± 0.89 μm), and 4-days post-pMCAO (WT: 39.24 ± 1.88 μm; KO: 47.43 ± 1.87 μm) (Figure 6A-6B). This was also reflected across a range of diameter distributions, showing an increased percentage of KO collaterals greater than 40µm (Supplementary Figure 5C-5F). Conversely, no differences were seen between WT and KO MCA-PCA collateral vessels at any time point (Supplementary Figure 5H).

**Figure 6.**
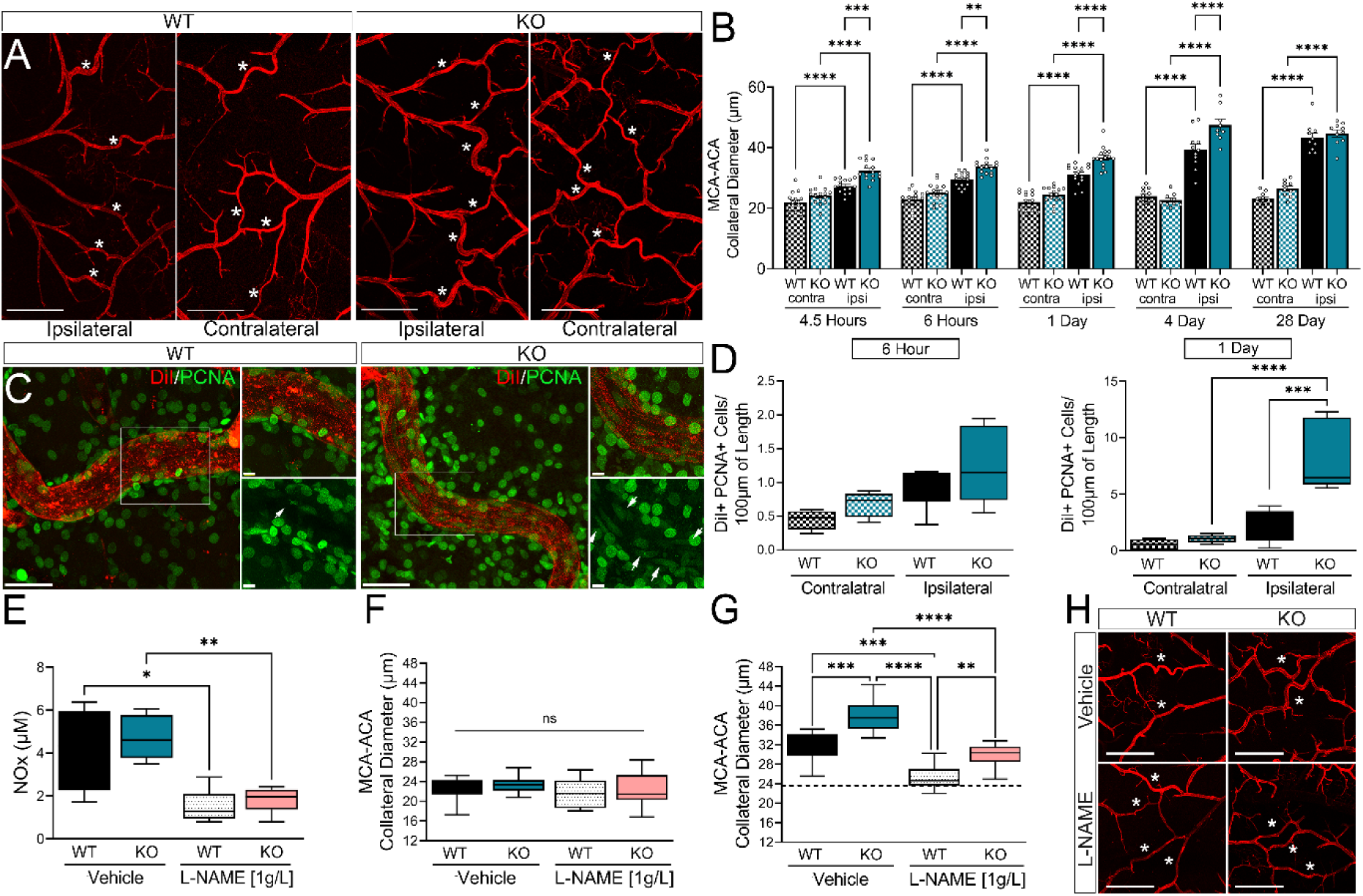
EC-specific EphA4 KO improves collateral growth and is partially attenuated with L-NAME treatment after pMCAO. (A) Representative 4X tiled images of the MCA-ACA pial collateral niche in WT and KO mice. (B) Analysis of MCA-ACA collateral diameter shows increased vessel size from 4.5-hrs to 4-days post-pMCAO in KO mice compared to WT controls. No difference is seen at 28-days post-stroke. n=9-15 Scale Bar = 1mm. (C) Representative 40X and 60X images of PCNA stained WT and KO MCA-ACA collaterals at 24hrs. (D) Quantification of PCNA-positive, DiI vessel painted collaterals shows increased proliferation at 24 but not 6hrs in KO mice. n=5. Scale Bar = 40µm for 40X images and 10µm for 60X. (E) Mice treated with 1g/L of L-NAME in drinking water shows reduced serum NOx levels at 24hrs compared to control mice. n=4-5. (F) No difference was observed in contralateral MCA-ACA collateral diameter in mice treated with vehicle or L-NAME. (G) In the ipsilateral hemisphere, L-NAME treatment significantly decreased collateral diameter in both WT and KO mice compared to vehicle controls. However, ipsilateral collateral diameter remained elevated in L-NAME KO compared WT mice. (H) Representative 4X images of ipsilateral MCA-ACA collateral vessels. n=9-11. Scale Bar=500 µm. Analysis was done using two-way ANOVA with Tukey’s post-hoc test. Dashed line in panel G = average contralateral collateral diameter. *p<0.05, **p<0.01, ***p<0.001, ****p<0.0001.

To uncover the nature of the cellular changes in the MCA-ACA collateral vessels that result in the increased diameter in KO mice, immunolabeling was performed on vessel-painted whole mounts of the dorsal cortex from WT and KO mice following pMCAO. Analysis of CD11b and Iba1 staining showed no difference at 4.5 hours between WT and KO mice. However, by 6hrs, KO mice show significantly higher total CD11b+ immune cell recruitment than WT controls (WT: 0.95 ± 0.21 cells/100μm; KO: 2.37 ± 0.35 cells/100μm) (Supplementary Figure 6A-6B). KO mice also display higher recruitment of CD11b+/Iba1+ monocytes/macrophages (WT: 0.71 ± 0.14 cells/100μm; KO: 1.63 ± 0.32 cells/100μm) and CD11b+/Iba1-immune cells compared to WT (WT: 0.24 ± 0.08 cells/100μm; KO: 0.73 ± 0.15 cells/100μm) (Supplementary Figure 6C). At 1-day post-pMCAO, KO mice maintain significantly higher total CD11b+ immune cell recruitment to collateral vessels (Supplementary Fig. 6B; WT: 1.80 ± 0.19 cells/100μm; KO: 3.1 ± 0.61 cells/100μm), but no differences are seen between genotype in immune subtype recruited. (Supplementary Figure 6B-6C).

Similar to Vasculotide-treated mice, EphA4 KO mice exhibited increased PCNA+/DiI+ proliferating ECs at 1-day. This increase was not present at 6hrs post-pMCAO (Figure 6C-6D). Moreover, these changes are not associated with alterations in smooth muscle cell (SMC) coverage or reorganization (Supplementary Figure 6D-6F), suggesting immune cell recruitment and endothelial cell division may provide key remodeling changes that support the acute acceleration of collateral growth in the absence of EC-specific EphA4 after ischemic stroke.

Lastly, NO-induced vasodilation was evaluated in the EphA4 KO mice 1-day post-pMCAO using L-NAME. A significant reduction in serum metabolites of NO, nitrate/nitrite (NOx) levels was observed in WT and KO mice, confirming treatment regime efficacy (Figure 6E). While no difference was seen in the contralateral hemisphere (Vehicle: 22.34 ± 0.75 μm vs. 23.45 ± 0.58 μm; L-NAME: 21.63 ± 0.95 μm vs. 22.28 ± 0.97 μm), ipsilateral collateral size was attenuated in L-NAME-treated WT (L-NAME: 25.23 ± 0.78 μm vs. Vehicle: 29.95 ± 0.67 μm) and KO mice (L-NAME: 31.48 ± 0.91 μm vs. Vehicle: 37.69 ± 1.19 μm) at 1-day post-pMCAO. However, the ipsilateral L-NAME-treated KO collateral vessels remained significantly larger than L-NAME-treated WT vessels, suggesting additional factors may influence pMCAO-induced collateral growth (Figure 6F-6H).

### Meningeal transcriptomic reveals common alterations in the absence of EC-specific EphA4 after pMCAO

Bulk RNA sequencing was performed on the meninges in the presence or absence of EC-specific EphA4. Comparative RNA sequencing analysis of ipsilateral samples in WT and KO mice show significant changes in gene expression across conditions (Figure 7A), including increased Krt5 in KO mice that is similarly seen in Vasculotide-treated mice. Venn diagram outlines those changes (Figure 7B), including top 10 genes in between each condition (Figure 7D-7E). GO terms for genes altered in KO compared to WT ipsilateral tissue include cell adhesion, cell chemotaxis, adhesion, long-chain fatty acid metabolic process (Figure 7F). Between Vasculotide and KO mice compared to WT, changes seen included 2004 upregulated genes, and 1240 downregulated genes (Figure 7G, 7H) shared between all groups compared to contralateral samples. Mice treated with Vasculotide had the highest number of unique, differentially expressed genes, with 466 upregulated and 714 downregulated, compared to vehicle-treated WT or KO mice. Interestingly, there were 203 upregulated and 212 downregulated genes, including *Krt5, Krt14, Col17a1,* and *Pgam1-ps1* (Figure 7I-7J; Supplemental Table 1), that were shared between EC-specific KO and Vasculotide-treated mice, indicating mutual pathways underlying endothelial EphA4/Tie2 axis following ischemic stroke. This includes Krt5 expression, which is upregulated on arterial vascular network in KO mice. This represents a new and novel protein target for meningeal vascular response to stroke.

**Figure 7.**
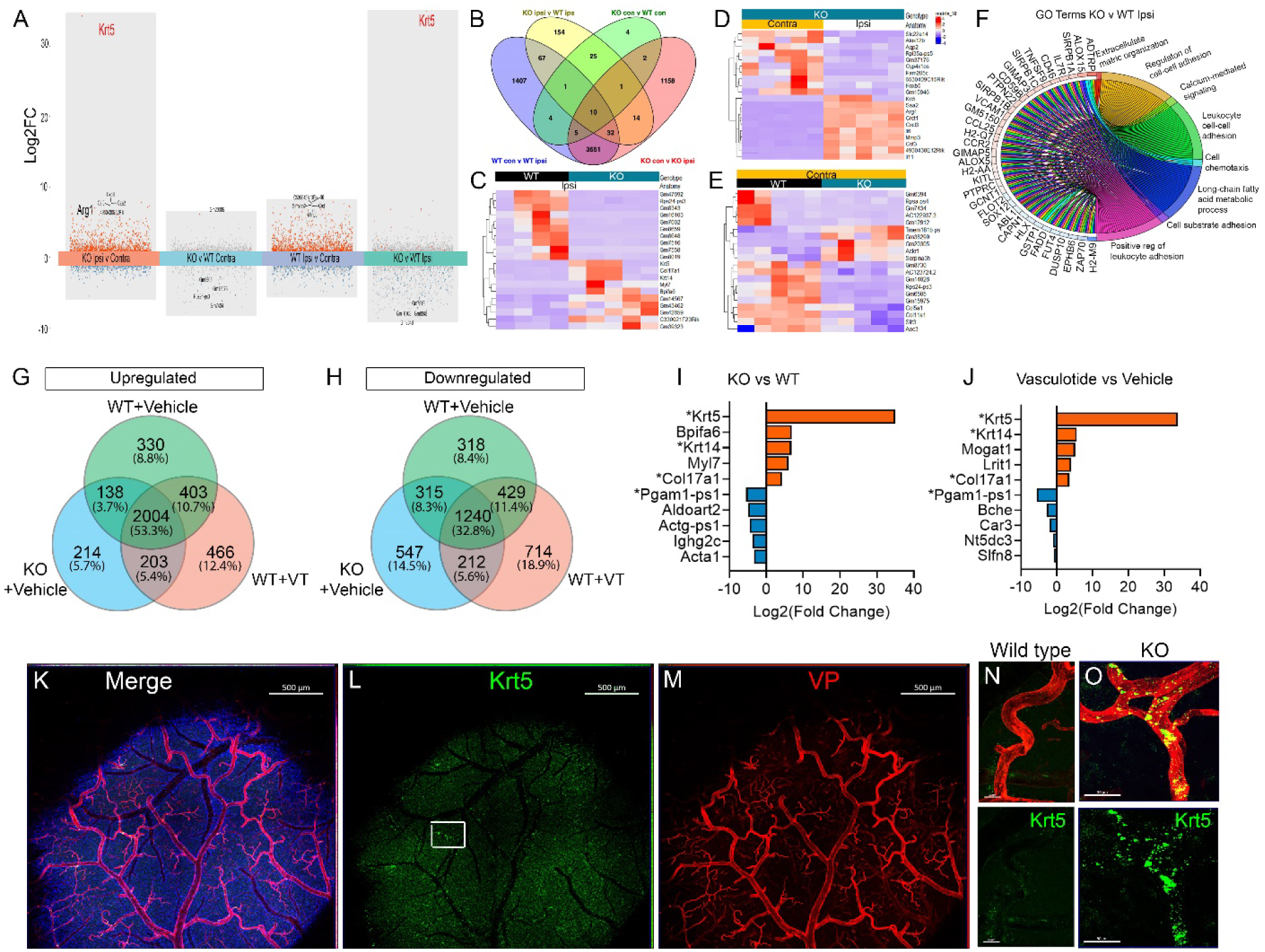
RNA sequencing of meningeal isolates displays altered transcriptomics in EphA4 mice that converges with Vasculotide-treated mice. (A) Transformed plot of bulk RNA sequencing from WT and KO meninges at 24hrs post-pMCAO. (B) Venn diagrams of upregulated and downregulated gene expression changes in ipsilateral vs contralateral pial surfaces of WT and KO mice 24hrs post-stroke. (D-E) Heatmap of top 10 upregulated and downregulated genes across conditions. (F) Chord plot of genes and Go terms between KO and WT ipsilateral meninges. (G-H) Venn diagram of up and downregulated genes between WT, KO and Vasculotide-treated conditions. (I-J) Common top 5 genes between conditions. (K-O) IHC of whole mounts showing increased Krt5 in KO arterial vasculature. Scale bar=500um in K-M and 50um in N and O.

## Discussion

The significance of pial collateral vessels in restoring blood flow to affected brain regions and shaping patient outcomes after ischemic stroke is well established. However, a notable lack of research focused on understanding the acute remodeling events and the critical mechanisms involved remains. The present study examined the temporal remodeling of pial collateral vessels and dissected the role of Tie2 and its regulators on collateral growth. Administration of a single dose of Tie2 agonist, Vasculotide, led to neuroprotection and improved functional outcomes, accompanied by increased MCA-ACA collateral size and cellular remodeling within the first 24 hours post-pMCAO. We also show that collateral growth and neuroprotection were attenuated in double Tie2 and EphA4 EC-specific knockout mice, demonstrating that EphA4 restricts collateral vessel growth by suppressing Tie2 signaling. Further analysis of the upstream regulator, EphA4, showed that EC-specific loss could mimic the effects of Vasculotide. In the EphA4 KO strain, we observed an increase in collateral size as early as 4.5 hours and substantially increased immune cell recruitment and proliferation in the vessel wall, suggesting these cellular changes contribute to the remodeling process. Interestingly, nitric oxide (NO) inhibition attenuated collateral growth in WT but only partially in Vasculotide-treated or EphA4 KO mice. This suggests that additional changes to collateral vessels may support growth and remodeling.

Tie2 is a receptor tyrosine kinase that plays a pivotal role in vascular development, with its disruption being embryonically lethal[14]. Ang-1/Tie2 signaling plays a dual role in vascular maintenance and growth in adulthood. Prior work established the Tie2 antagonist, angiopoietin-2 (Ang-2), suppresses collateral growth in hindlimb ischemia[9]. We show that a single treatment of Vasculotide can enhance arteriogenesis, the first evidence that the Ang-1 peptide agonist can influence collateral vessels in ischemic stroke. Additional studies have shown that treatment with 3μg/kg Vasculotide delivered i.p. in diabetic rats subjected to stroke showed decreased infarct volume and improved functional recovery[15]. Vasculotide also improved white matter recovery and increased vascular density at the ischemic border when delivery is delayed 24 hours post-stroke and given daily for 14 days[16]. This suggests that Vasculotide influences both acute and subchronic remodeling and repair responses.

Our work with Tie2 and double EphA4/Tie2 KD mice demonstrates that loss of EC-specific EphA4 mediates collateral growth by enhancing Tie2 signaling. This effect may be due to increased expression of Tie2, altered ligand activation, or downstream signaling such as p-Akt[17]. In adulthood, EphA4 has been implicated in numerous disease and injury processes. Previous work in stroke models has shown that EphA4 activation exacerbates brain edema and reduces functional recovery[18] while inhibiting EphA4 could confer neuroprotection and enhance collateral response[11]. Furthermore, the use of Tie2-Fc in EphA4 KO mice to block Tie2 signaling attenuated their response back to WT levels, further supporting the connection between EphA4 and Tie2[11]

Transcriptomic alterations were observed in the meninges of both EC-specific EphA4 KO and Vasculotide-treated mice, which uniquely shared a key change in *Krt5, Krt14, and Col17a1* expression. These genes are markers for epidermal stem cells [19]. They are key components of desmosomes and adherens junctions (AJ), which mediate cell-cell contact sites and aid in integrating chemical and mechanical signaling [20]. Loss of function studies on AJ components have relied on targeted gene ablation using the keratin 5 or 14 promoter [21]. Previous findings show increased organization of AJ and diameter expansion of coronary collateral reserve [22]. It remains unclear whether pial collateral growth may be further influenced by AJ remodeling. Future studies may reveal whether modulating *Krt5* could promote this avenue of collateral expansion following ischemic stroke.

Future work should continue to evaluate the efficacy of Vasculotide as a therapeutic enhancer for pial collateral function after ischemic stroke and aim to elucidate how a single dose of the drug can result in sustained improvements. Additionally, while there is limited evidence to show that Tie2 displays any sex differences in its baseline levels or function, the efficacy of Vasculotide for enhancing collateral growth should be investigated in females as stroke outcomes can differ by sex. Lastly, since ischemic stroke in humans often occurs alongside comorbidities like hypertension, future studies should assess Vasculotide therapeutic potential in mouse models with this clinical condition. This will provide an additional clinically relevant understanding of the peptide’s efficacy and help guide its translation to human trials.

In conclusion, our findings provide evidence of crosstalk between EphA4 and Tie2 in endothelial cells and offer insights into the acute cellular changes in pial collateral vessels after pMCAO. Understanding the mechanisms of collateral growth is crucial for developing targeted therapeutics to enhance this adaptive response, and our results indicate that activation of Tie2 signaling represents a promising novel therapeutic strategy.

## Methods

### Animals

Mice were housed in a virus/antigen-free, AALAC-accredited facility on a 12h light/dark cycle with standard rodent chow and water ad libitum. Male mice were used for experiments at 8-12 weeks of age. For Vasculotide experiments, male CD1 mice were utilized. *<u>Sex as a biological variable:</u>* Male mice were selected for these experiments to minimize variability introduced by hormonal fluctuations inherent to female mice during the estrous cycle, which could impact vascular and immune responses, which may potentially mask or alter the interpretation of our findings. By using male mice, we aim to reduce biological variability, thereby improving the reproducibility and interpretability of the results. Additionally, male mice are commonly used in similar studies, allowing for better comparability with existing literature. Future studies will include both sexes to determine whether the observed mechanisms and outcomes are sex-specific. EphA4 knockout experiments used *EphA4^fl/fl^* (WT) and *EphA4^fl/fl^/VECadherin-Cre^ERT2^* (KO) were bred on CD1 background for at least 10 backcrosses. For Tie2 genetic deletion experiments, *Tie2^+/+^/EphA4^+/+^/VECahherin-Cre^ERT2^*(WT), *Tie2^fl/fl^/EphA4^+/+^/VECaherin-Cre^ERT2^*(tKD), *Tie2^+/+^/EphA4^fl/fl^/VECaherin-Cre^ERT2^*(eKD), and double knockdown mice *Tie2^fl/fl^/EphA4^fl/fl^/VECaherin-Cre^ERT2^* (dKD) were generated on a mixed CD1 and C57BL/6 background. Each animal was coded, and the experimenter was blinded from group conditions.

### Tamoxifen Injections

Mice were intraperitoneally injected for five consecutive days starting at eight weeks of age with 2mg/mouse or 50mg/kg of tamoxifen (Sigma Aldrich; St Louis, MO, USA) diluted in corn oil. Two weeks following last injection, tail snips were taken for genotyping, as previously described [17].

### Vessel Painting and collateral quantification

Vessel painting was performed as previously described [11, 23]. Whole brain tiled images were taken at 4X magnification using a Nikon C2 confocal (Tokyo, Japan). Fiji-ImageJ (NIH) version 2.14.0 was used for quantification.

### Surgical Procedures and Treatments

Ischemic stroke was induced in adult mice via permanent middle cerebral artery occlusion, as previously described [11]. Briefly, mice were anesthetized with 2.5%isolfulane-30% oxygen and received Buprenorphine-SR (3.25mg/kg; EthiqaXR, Fidelis Animal Health, USA) and the main distal branch and two bifurcating branches were cauterized. Sham mice received the same procedure, excluding ligation. For Vasculotide experiments as previously described[17], mice randomly received either saline (vehicle control), 3μg/kg Vasculotide, or 150μg/kg Vasculotide via tail vein bolus injection immediately following sham or pMCAO. For clustered (cl) EphA4-Fc experiments, mice received either clustered 1mg/kg clEphA4-Fc or 0.34mg/kg of soluble human Fc control via tail vein bolus injection immediately following pMCAO. Mice were euthanized by vessel painting between 4.5 hours and 28 days post-surgery. A total of 556 mice were used, with the overall mortality rate of the procedure being 2.34% (13 of 556 mice). As a result, 543 were included in the data presented within this manuscript.

### Nitric Oxide Synthase Inhibitor Treatment

Following pMCAO surgery, mice were randomly placed on L-NAME dissolved in drinking water (1g/L) as previously described [24]. Mice were allowed ad libitum access for 24hrs, at which point they were euthanized by vessel painting or for serum collected. Serum samples were processed following manufacturer guidelines for a nitrate/nitrite colorimetric analysis kit (Cayman Chemical, Ann Arbor, MI, USA) to quantify the end products of nitric oxide metabolism, nitrate (NO_3_^-^) and nitrite (NO_2_^-^).

### Behavioral Testing

All animal behavioral assessments were performed as previously described [11, 25, 26]. *Rotarod.* Mice were trained for 4 consecutive days before pMCAO or sham surgery, with baseline measurements recorded on day four and testing performed on days 3-28. Four trials were performed per day at 4 rpm and an acceleration of 0.1rpm/sec, with two minutes of rest between trials. *Modified Neurological severity scoring (mNSS).* Deficit was graded on a 0-14 scale (0=normal function; 14=maximum deficit) based on motor, balance, and reflex tests. Baseline measurements were taken one day before pMCAO or sham surgeries and mice were tested on days 1, 3, 7, 14, 21, and 28 following the procedure. *Adhesive Tape Test*. A piece of 3mm x 4mm cloth adhesive tape was placed on each forepaw with equal pressure. The mice were returned to a transparent testing box, where the time-to-contact and the time-to-remove were recorded for the ipsilateral and contralateral forepaws. Before sham or pMCAO surgery, mice underwent four consecutive days of training and then tested on days 1, 3, 7, 14, 21, and 28 following surgery. Scores were reported as asymmetry scores, as previously published [27]. Asymmetry Score = (contralateral time to remove – ipsilateral time to remove)/(contralateral time to remove + ipsilateral time to remove)*100.

### Cerebral blood flow (CBF)

Blood flow analysis was done using laser speckle contrast imaging (RFLSI III Laser Speckle Imaging System, RWD, China) through a thinned skull. Briefly, mice were anesthetized with 2.5% isoflurane, ventilated with a mixture of O2 and air (30:70), the skin was prepared, and a midline incision was made. A drill with a carbide bur was used to thin the skull as previously described [28]. The mice were placed in a stereotactic adaptor under the LSCI system with a working distance of 15cm, and warmed saline was continuously applied to the thinned skull to keep it moist. Temperature was monitored at 37 ± 0.5°C, and imaging was performed over 60 seconds using the HD algorithm. CBF was assessed pre-injury and 10m, 6hrs, and 1-4 days post-injury. A standardized region of interest (ROI) was used for each mouse, and perfusion units were calculated relative to the contralateral, uninjured hemisphere.

### Infarct volume

Serial coronal sections cut at 30μm slices and 990μm apart were stained with 0.2% Cresyl violet (Electron Microscopy Science, Hatfield, PA, USA). The loss of Nissl staining on six serial sections was quantified using the Cavalieri Estimator, StereoInvestigator software (MicroBrightField, Williston, VT, USA), as previously described [11, 29].

### Immunohistochemistry of cortical whole mounts

Whole cortical mounts were dissected, washed in 1xPBS, and placed in 2% Fish Gel with 0.4% Triton block, then incubated for two days in rabbit anti-smooth muscle actin (SMA) (Cell Signaling Technology, Danvers, MA, USA; 19245S), rat anti-CD11b (Abcam, Cambridge, UK; ab8878), or rabbit anti-Iba1 (FUJIFILM WakoPure Chemical Corporation, Osaka, Japan; 019-19741) primary antibodies in block at 1:200. Following washing with 1XPBS with 0.1% Tween 20, whole cortical samples were incubated for two hours at room temperature with Alexa Flour 488 or Alexa Fluor 647 conjugated secondary antibodies (ThermoFisher, Waltham, MA). Samples were washed with 1XPBS + 0.1% Tween 20 and imaged on a Nikon C2 inverted confocal microscope (Tokyo, Japan). For PCNA staining, samples were first incubated for 1.5hrs in 6N HCl at 37°C, followed by a 30-minute incubation in sodium borate to neutralize the acid treatment before using rabbit anti-PCNA (1:300; Cell Signaling Technology, Danvers, MA, USA; 13110).

### Endothelial Cell Culture

Primary, brain-derived murine endothelial cells were isolated, characterized, and cultured [10]. For western blot analysis, ECs were treated in base media with 10nM Vasculotide or PBS, incubated for 5 minutes at 37C, washed, and resuspended in RIPA buffer containing proteinase and phosphatase inhibitors.

### Western Blot analysis

Tissue or cells were homogenized on ice in RIPA buffer, and homogenates were spun at 4°C for 20 minutes at 15,000 x g, then stored at -80°C until use. Protein concentration was determined using the PierceTM BCA Protein Assay Kit (ThermoFisher, Waltham, MA, USA), and fifty micrograms of protein per sample were run on an 8% gel for Tie2/pTie2 or 10% gel for Ang1/Ang2, then transferred onto a PVDF membrane. Membranes were blocked overnight using EveryBlot blocking buffer (Biorad, Hercules, CA, USA) then incubated at 4°C with primary antibodies, pTie2 (ThermoFisher, Waltham, MA, USA), Tie2, Ang1(R&D Systems, Minneapolis, MN, USA), Ang-2 (Abcam, Cambridge, UK), and β-actin (Cell Signaling Technology, Danvers, MA, USA). Following incubation, membranes were washed in 1XTBST, incubated for 1hr in fluorescent secondary antibodies 1:5000 (LI-COR Biosciences, Lincoln, NE, USA), and imaging was performed on Odyssey Imaging System (LI-COR, Lincoln, NE, USA), and relative density analyzed using Fiji-ImageJ (NIH).

### Genome-wide RNA bulk Sequencing

Based on previous literature[3, 30], mice were cardiac perfused with 0.9ml of RNAlater (Sigma Aldrich) with 0.1ml 2% Evans blue dye, and brains were frozen overnight in RNAlater and then pial surfaces removed the following day. Samples were pooled together from four mice and placed in TRIzol® reagent (ThermoFisher, Waltham, MA. USA). RNA was isolated using the Direct-zol RNA microprep kit (Zymo, Irvine, CA) and total RNA from the pial surfaces according to manufactures instructions and was quantified by absorbance with spectrophotometer ND-1000 (ThermoFisher, Waltham, MA. USA). 1μg total RNA was used for RNA-seq library construction, MedGenome Inc. (Foster City, CA, USA). The libraries were sequenced on Novaseq platform with 100 bp paired-end mode (Illumina). Data quality check was performed using FastQC (v0.11.8). The adapter trimming was performed using fastq-mcf program (v1.05) and cutadapt (v2.5). For the RNA-Seq analysis, we remove the unwanted sequences, including mitochondrial genome sequences, ribosomal RNAs, transfer RNAs, adapter sequences, and others. Contamination removal was performed using Bowtie2 (v2.5.1). Clean reads were mapped to the GRCm38.p6 genome, and Alignment was performed using STAR (v2.7.3a) aligner. Reads mapping to ribosomal and mitochondrial genomes were removed before performing alignment. The raw read counts were estimated using HTSeq (v0.11.2). R package DESeq2 was applied to get the normalized counts and downstream analysis. Genes with larger than 1.2-fold changes and adjusted p-values of less than 0.05 were considered significant.

### Statistical analysis

The data was analyzed and plotted with GraphPad Prism, version 9 (GraphPad Software, Inc., San Diego, CA, USA). To compare two experimental groups, the Student’s two-tailed t-test was employed. For multiple comparisons, one-way or two-way ANOVA and repeated measures were used, followed by post hoc Tukey’s analysis where appropriate. Changes were considered significant at P< 0.05. Mean values, accompanied by the standard error of the mean (SEM), were reported.

### Study Approval

All experiments were performed under the NIH Guide for the Care and Use of Laboratory Animals and under the approval of the Virginia Tech Institutional Animal Care and Use Committee (IACUC; #21-056).

### Data Availability

All sequencing data can be found in the NIH Gene Expression Omnibus under code: GSE242163.

### Author Contribution

A.M.K., K.L., N.A.G., Y.L., H.X., and M.H.T. performed research and analyzed data. A.M.K., J.Z., H.X., J.B.M., and M.H.T contributed reagents/analytic tools. A.M.K. and M.H.T. designed the research and wrote and edited the paper.

## Acknowledgments

We recognize the Biomedical and Veterinary Sciences Program (BMVS). We thank Dr. John Chappell for the generous gift of *VeCadherin-CreERT2* mice and Dr. Hellmut Augustin for the gift of *Tie2^fl/fl^* mice. We recognize Xiguang Xu and the Fralin Life Sciences Genomic Sequencing Center for assistance with RNA quality testing. This work was supported by the National Institute of Neurological Disorders and Stroke of the National Institutes of Health, R01NS112541 (MHT).

## Non-Standard Abbreviations and Acronyms

WT: wildtype
KO: knockout
tKD: Tie2 knockdown
dKD: double Tie2/EphA4 knockdown
pMCAO: permanent middle cerebral artery occlusion
MCA: middle cerebral artery
ACA: anterior cerebral artery
PCA: posterior cerebral artery
CBF: cerebral blood flow
EC: endothelial cells
SMC: smooth muscle cell
SMA: smooth muscle actin
Ang: angiopoietin
VT: Vasculotide

---

**Supplemental Figure 1.**
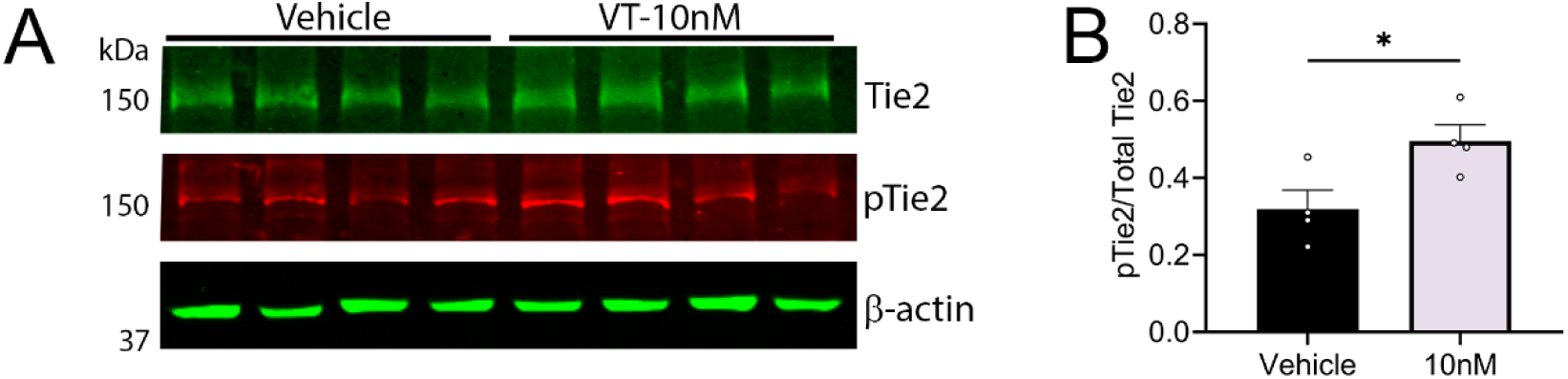
Tie2 peptide agonist Vasculotide (VT) stimulates pTie2 *in vitro*. (A) Western blot analysis of wild-type primary endothelial cells after treatment with 10nM Vasculotide or vehicle for 5 minutes. (B) Quantification of pTie2 normalized to Tie2 expression reveals increased pTie2 in Vasculotide-treated cells. N=4; *P<0.05.

**Supplemental Figure 2.**
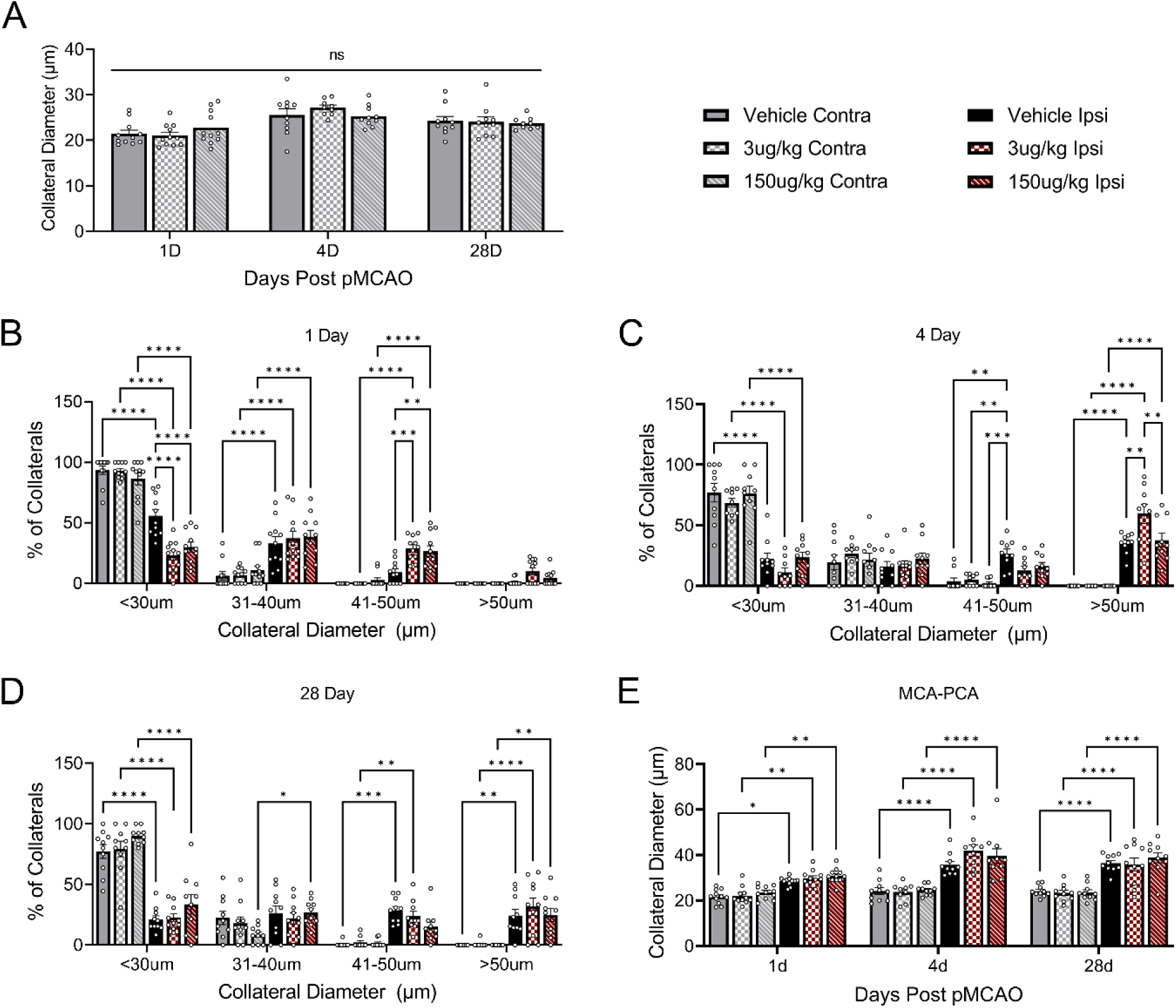
Vasculotide alters ipsilateral but not contralateral MCA-PCA collateral size. (A) Independent of dose, Vasculotide did not alter collateral size in the contralateral hemisphere post-pMCAO. (B) Vasculotide treatment resulted in fewer MCA-ACA collaterals under 30 µm and more in the 41-50 µm range at 1 day post-surgery. (C) By 4 days post-pMCAO, only the 3ug/kg Vasculotide treatment displayed more collaterals in the greater than 50 µm range. (D) No difference was found in MCA-ACA collateral distribution at 28 days post-stroke. (E) Vasculotide treatment in the MCA-PCA collateral niche did not alter collateral size at any time point. N=9-11 mice. Analysis was done using two-way ANOVA with Tukey’s post-hoc test. *p<0.05, **p<0.01, ***p<0.001, ****p<0.0001.

**Supplemental Figure 3.**
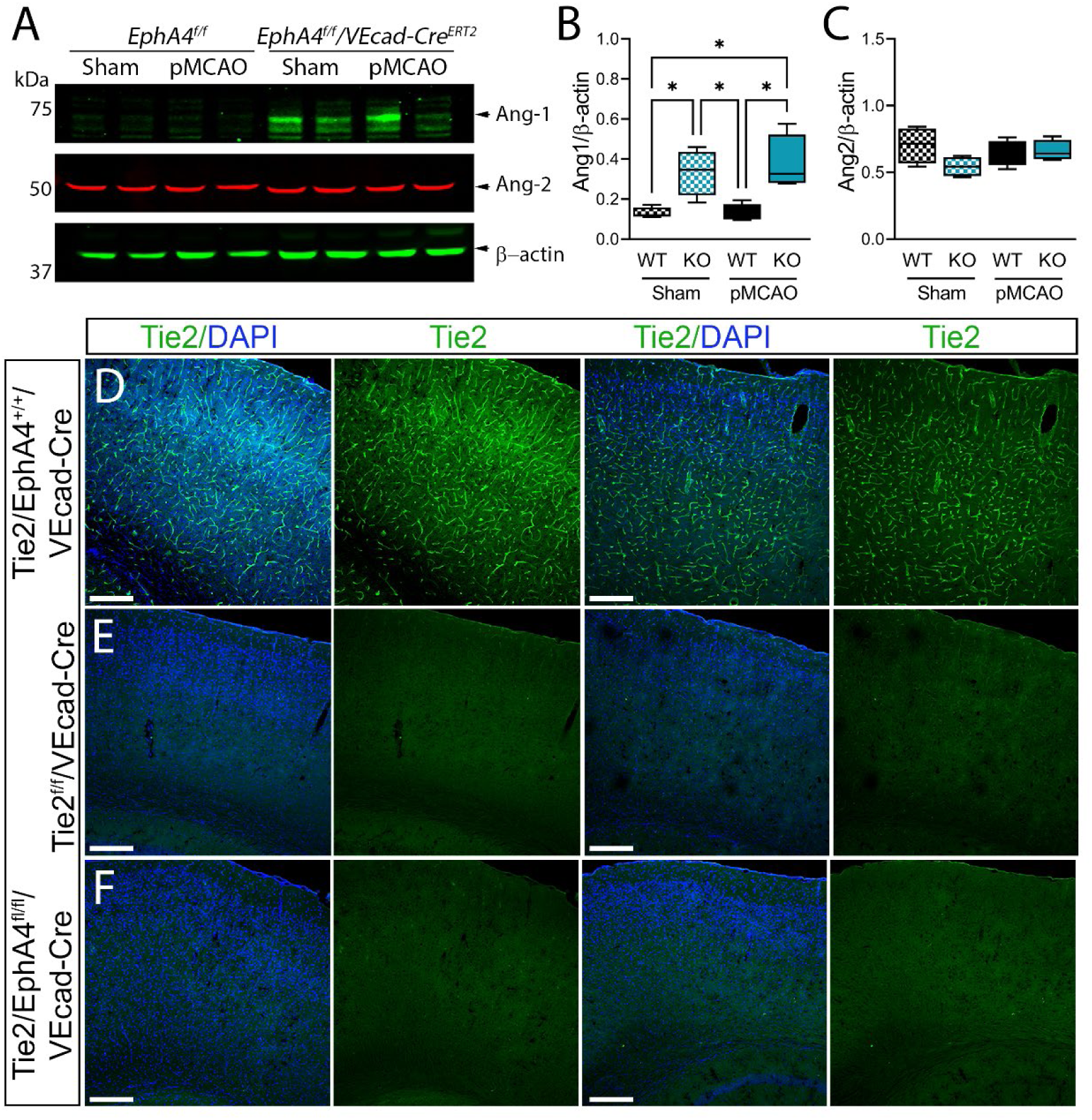
Tie2 expression is eliminated in tKD (*Tie2^fl/fl^/VECaherin-Cre^ERT2^*) and dKD (*Tie2^fl/fl^EphA4^fl/fl^/VECaherin-Cre^ERT2^*) mouse lines. (A) Western blot analysis of WT and EphA4 KO cortex samples 24hrs post-pMCAO. (B) Densitometric analysis shows a marked increase in Ang-1 in the EphA4 KO cortex compared to WT after stroke. (C) No difference was observed in Ang-2 expression. N=4. (D) Representative 10X confocal images of Tie2 stained serial sections revealed notable expression in Cre-positive wildtype mice. Loss of staining was observed in (E) tKD and (F) dKD mice. Scale = 100um. Analysis was done using a two-way ANOVA with Tukey’s post-hoc test. *p<0.05, **p<0.01, ***p<0.001.

**Supplemental Figure 4.**
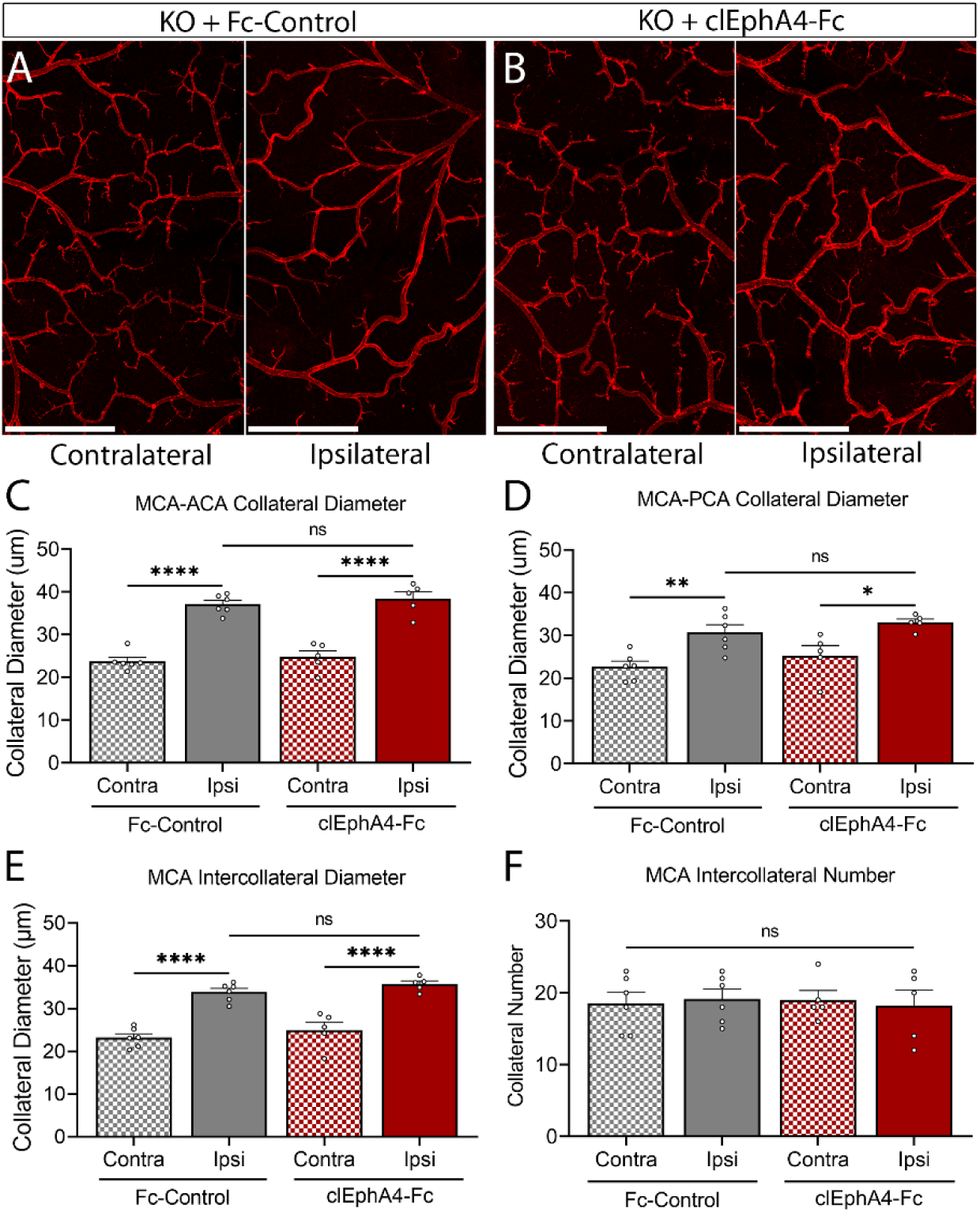
EC EphA4 forward signaling regulates collateral growth after pMCAO. (A) Representative 4X tiled confocal images of vessel-painted MCA-ACA pial collaterals in KO mice that received either clustered (cl) Fc-control or (B) clEphA4-Fc recombinant proteins directly following pMCAO. (C) No differences were observed between the ipsilateral collateral size of KO mice that received clEphA4-Fc or clustered Fc-control in the MCA-ACA or (D) MCA-PCA pial collaterals. (E) Neither MCA intercollateral diameter nor (F) number was altered by treatment with clEphA4-Fc compared to Fc-control treated mice. N=5-6 mice. Scale bar = 1mm. Analysis was done using a two-way ANOVA with Tukey’s post-hoc test. *p<0.05, **p<0.01, ****p<0.0001

**Supplemental Figure 5.**
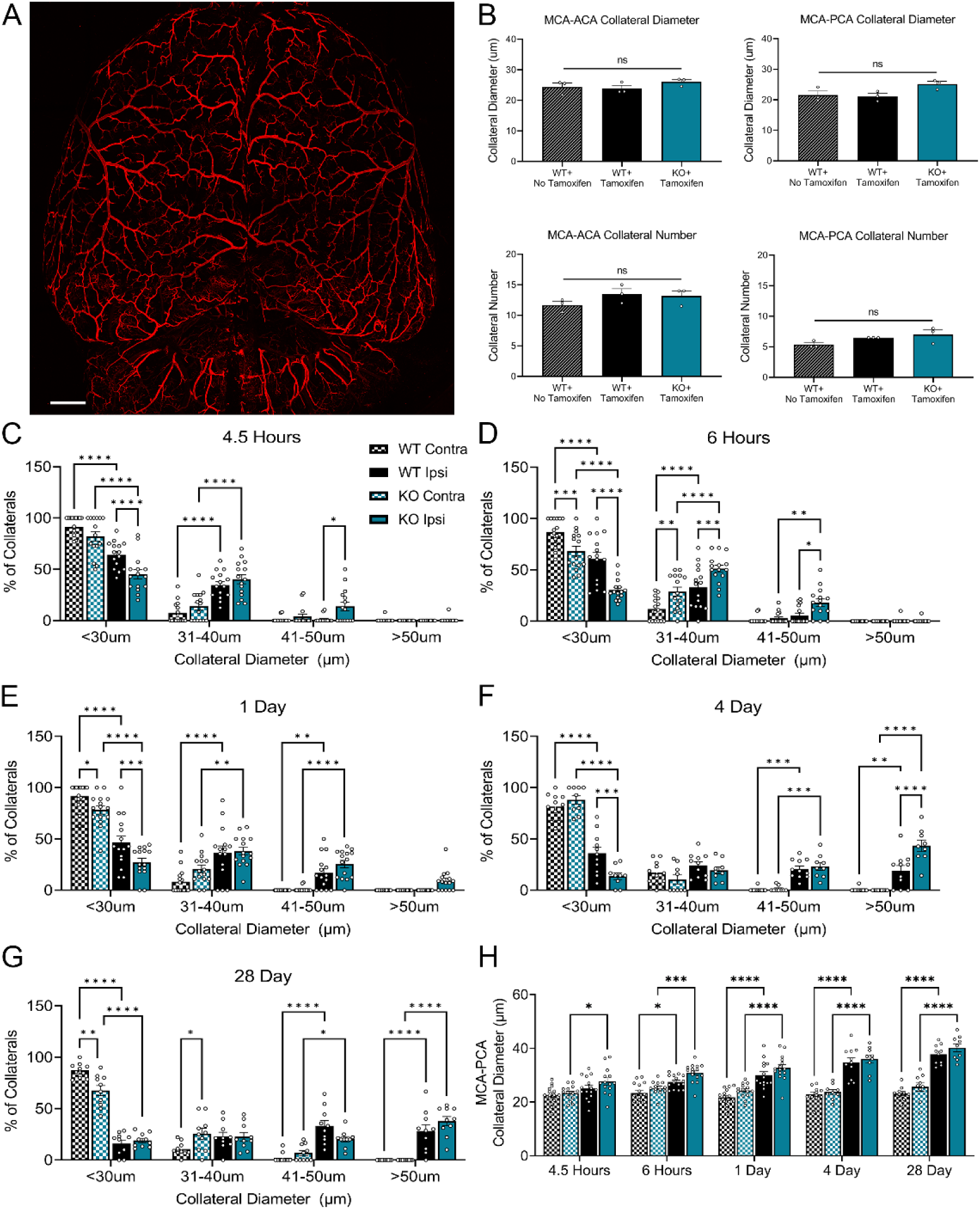
EC-specific loss of EphA4 does not influence collateral size or number in naïve mice. Naïve mice were vessel painted to assess how the removal of EphA4 from endothelial cells affected collateral size. (A) Representative 4X tiled confocal image of a naïve vessel painted brain. (B) Loss of EphA4 did not impact collateral size in either the MCA-ACA (top left) or MCA-PCA (top right) collateral niches. The collateral number was not altered by tamoxifen injections (Bottom row). n=3. (C) Following pMCAO, the distribution of MCA-ACA connecting collaterals at 4.5hrs, (D) 6hrs, (E) 1 day, and (F) 4-days indicates KO mice have significantly fewer collaterals in the less than 30 µm and a higher percentage of collaterals in the 41-50 or >50-micron range compared to WT controls. (G) At 28 days post-pMCAO, no difference was seen in collateral distribution on the injured hemisphere by genotype. (H) No difference was noted in the collateral diameter of MCA-PCA connecting collaterals between WT and KO mice. n=9-15. Scale bar = 1mm. Analysis was done using a one-way (B) and two-way (C-H) ANOVA with Tukey’s post-hoc test. *p<0.05, **p<0.01, ***p<0.001, ****p<0.0001.

**Supplemental Figure 6.**
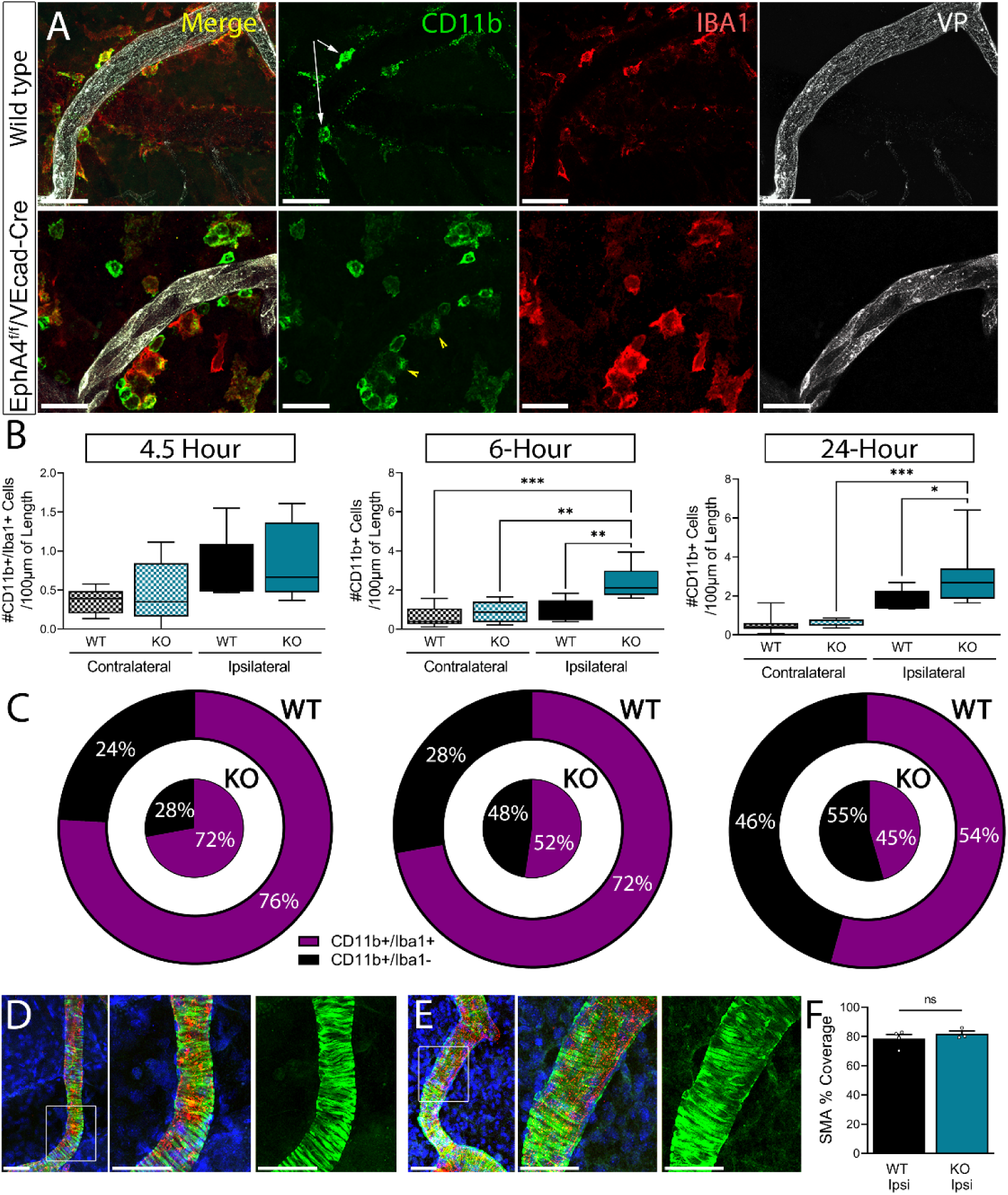
EphA4 ablation increases immune cell recruitment but fails to alter smooth muscle cell coverage following pMCAO. (A) Immune cell recruitment was quantified in cortical whole mounts stained with CD11b and Iba1. Representative images of WT and KO MCA-ACA pial collaterals at 24hrs post-pMCAO. (B) Analysis of total CD11b+ shows no change in immune cell recruitment between the genotypes at 4.5hrs but increases in recruitment to the KO ipsilateral collaterals at 6hrs and 24hrs post-pMCAO compared to WT collaterals. (C) Pie charts represent the percentage of immune cells recruited that are CD11b+/Iba1- (black) and CD11b+/Iba1+ (pink). N=5-7 mice. (D-E) Representative images of MCA-ACA connecting collaterals stained with SMA to show SMCs. (F) Analysis of percent coverage of SMA with DiI shows no significant change in SMC reorganization at 24 hours post-pMCAO. Analysis was done using two-way ANOVA with Tukey’s post-hoc test (B) or t-test (F). N=4-5 mice. Scale bar = 50um. p<0.05, **p<0.01, ***p<0.001, ****p<0.0001.

